# A low-footprint, fluorescence-based bacterial time-kill assay for estimating dose-dependent cell death dynamics

**DOI:** 10.1101/2024.03.08.584154

**Authors:** Eshan S. King, Anna E. Stacy, Jacob G. Scott

## Abstract

Dose-response curves that describe the relationship between antibiotic dose and growth rate in bacteria are commonly measured with optical density (OD) based assays. While being simple and high-throughput, any dose-dependent cell death dynamics are obscured, as OD assays in batch culture can only quantify a positive net change in cells. Time-kill experiments can be used to quantify cell death rates, but current techniques are extremely resource-intensive and may be biased by residual drug carried over into the quantification assay. Here, we report a novel, fluorescence-based time-kill assay leveraging resazurin as a viable cell count indicator. Our method improves upon previous techniques by greatly reducing the material cost and being robust to residual drug carry-over. We demonstrate our technique by quantifying a dose-response curve in *Escherichia coli* subject to cefotaxime, revealing dose-dependent death rates. We also show that our method is robust to extracellular debris and cell aggregation. Dose-response curves quantified with our method may provide a more accurate description of pathogen response to therapy, paving the way for more accurate integrated pharmacodynamic-pharmacokinetic studies.

## Introduction

Bacterial drug susceptibility is commonly quantified by minimum inhibitory concentration (MIC), which roughly summarizes the maximum drug concentration that a bacterial population can withstand. While MIC is a useful heuristic, especially in the clinic, dose-response curves provide a more complete description of pathogen response to a wide range of drug concentrations. For instance, dose-response curves, especially those that quantify growth rate as a function of drug concentration, can reveal fitness costs to drug resistance, where drug-resistant strains grow slower in the absence of drug compared to their drug-sensitive counterparts^1–4^. Connecting growth rate to drug concentration also permits more detailed models of pathogen response to clinical pharmacokinetics, where drug dosing and diffusion can generate highly variable drug concentrations within a patient^5–8^. Furthermore, the shape of antibiotic dose-response curves can provide insight into the biochemical mechanism of different drugs^3,9^.

Growth rate assays almost universally utilize optical density at 600 nm (OD_600_) over time to quantify the rate of change in cell count^1,2,4,9–11^. In some cases, OD_600_ is converted to cell count over time with a calibration curve, while others simply estimate growth rate from OD_600_ alone. When implemented with an automated microplate scanner, OD_600_ assays permit simple, high-throughput measurement of dose-response relationships. However, an important drawback of the OD assay in batch culture is that it only quantifies positive cell growth, and it cannot accurately or reliably detect cell death. Indeed, cell death simply appears as no change in optical density over time. Here, we use “net growth rate” to refer to the rate of change in cells over time. This term encapsulates both “birth rate”, or rate of cell division, and death rate into a single parameter:

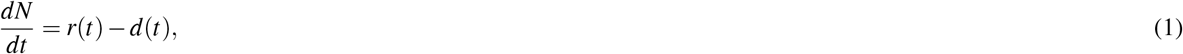

where *r*(*t*) is the birth rate, *d*(*t*) is the death rate, and 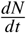 is the net growth rate. This drawback of the OD assay is highlighted in **Fig 1** – the assay can only measure a positive change in cells, and as such, the corresponding dose-response curve does not cross zero. Any dose-dependent cell death dynamics cannot be quantified in this way. In contrast, a time-kill assay quantifies *net* change in cells over time, including cell death, and a dose-response curve measured with a time-kill assay can cross zero and reveal dose-dependent cell death (**Fig 1B** and **1C**). Understanding dose-dependent cell death dynamics can better inform how pathogens respond to drug, allowing for more realistic integrated pharmacodynamic-pharmacokinetic (PK-PD) modeling of infections.

**Figure 1.**
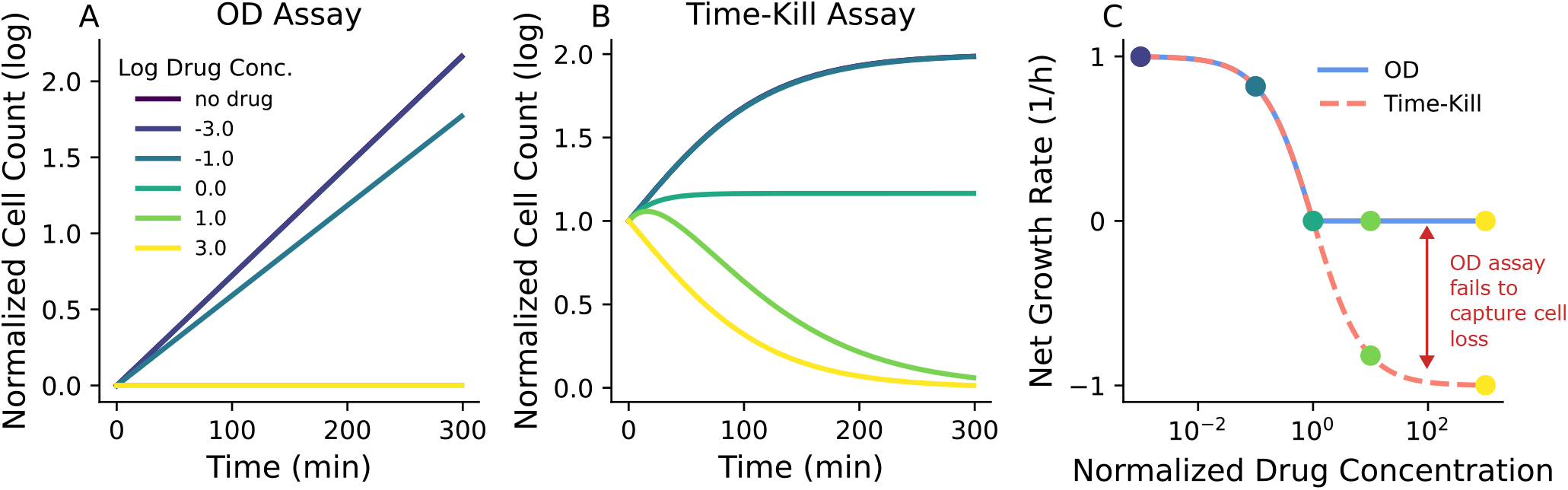
Traditional OD-based dose-response curves obscure killing dynamics. Illustrative dose-response experiments with synthetic data. **(A)** Cell count timeseries data estimated with optical density. Above the MIC (∼1 µg/mL), there is no detectable cell growth, and thus the curves are overlapping. **(B)** Cell count timeseries data from a time-kill assay experiment. (**C**) Dose-response curves estimated from the OD and time-kill assays. The OD assay fails to capture net growth rate less than 0. Colored data points correspond to the cell count timetraces in **A** and **B**.

Time-kill experiments have commonly been implemented with colony-forming unit (CFU) assays, where cultures are diluted and deposited onto agar plates such that single colonies can be counted and the average cell count can then be estimated^12–15^. Importantly, CFU assays are a direct estimate of cell count. However, the amount of time and resources required is a major drawback of the CFU method of time-kill assay. Each data point measured in triplicate requires 3 agar plates. In addition, it is normally unknown what the optimal dilution range is for accurate CFU counting, as too little dilution results in overlapping colonies while too much dilution results in unreliable or absent colony counts. Therefore, for each condition, a minimum of 2 plating dilutions are normally required, resulting in 6 agar plates for a single data point. To quantify a dose-response curve with a time-kill assay, one will need many time points to estimate the long- and short-term dynamics of the change in cell count in response to drug. As an example, 6 drug concentrations and 10 time points would require 360 agar plates for a single experiment, not including other materials. Furthermore, such an experiment is demanding for the researcher, requiring a high level of focus for many hours at a time, increasing the likelihood of mistakes.

Others have attempted to address these shortcomings by developing assays that rely on bacterial regrowth^16,17^. Briefly, small samples are periodically taken from a cell culture with drug and inoculated in fresh medium. Optical density is monitored over time, and the time it takes for a regrown culture to reach a specific OD threshold is then linearly related to the initial cell count, which can be calibrated with a CFU assay. While higher throughput than a traditional CFU time-kill assay, bacterial regrowth assays suffer from drug carry-over, where drug from the experimental culture is carried over into the regrowth culture. This residual drug may influence the regrowth dynamics and bias results. There is a need for a facile, low-footprint, and accurate method for estimating dose-dependent cell death dynamics in bacteria.

Resazurin, commercially marketed as Alamar Blue (AB), is a blue and weakly fluorescent dye that is rapidly reduced to the pink and highly fluorescent resofurin in the presence of viable cells^18,19^. Resazurin has been commonly used as a cell viability indicator in both prokaryotic^20,21^ and eukaryotic cells^18,22^. While resazurin has seen some use in bacterial cytotoxicity assays, it has not been used for full time-kill assays to quantify dose-dependent cell death rates^20,23,24^. Here, we present a novel time-kill assay utilizing resazurin to estimate viable cell count over time. An overview of the method is shown in **Fig 2**. Briefly, bacterial cell cultures are subjected to different drug concentrations in a 96-well plate. At designated timepoints, AB is added to wells and incubated for 30 minutes before scanning for fluorescence. The fluorescence value is then converted to cell count with a calibration curve. By repeating the process for several timepoints and multiple 96-well plates, one can estimate cell count dynamics under a range of different drug concentrations with multiple replicates in a single experiment. We use this method to quantify a pharmacodynamic relationship between the net change in *Escherichia coli* cells and, in this case, cefotaxime (CTX) concentration.

**Figure 2.**
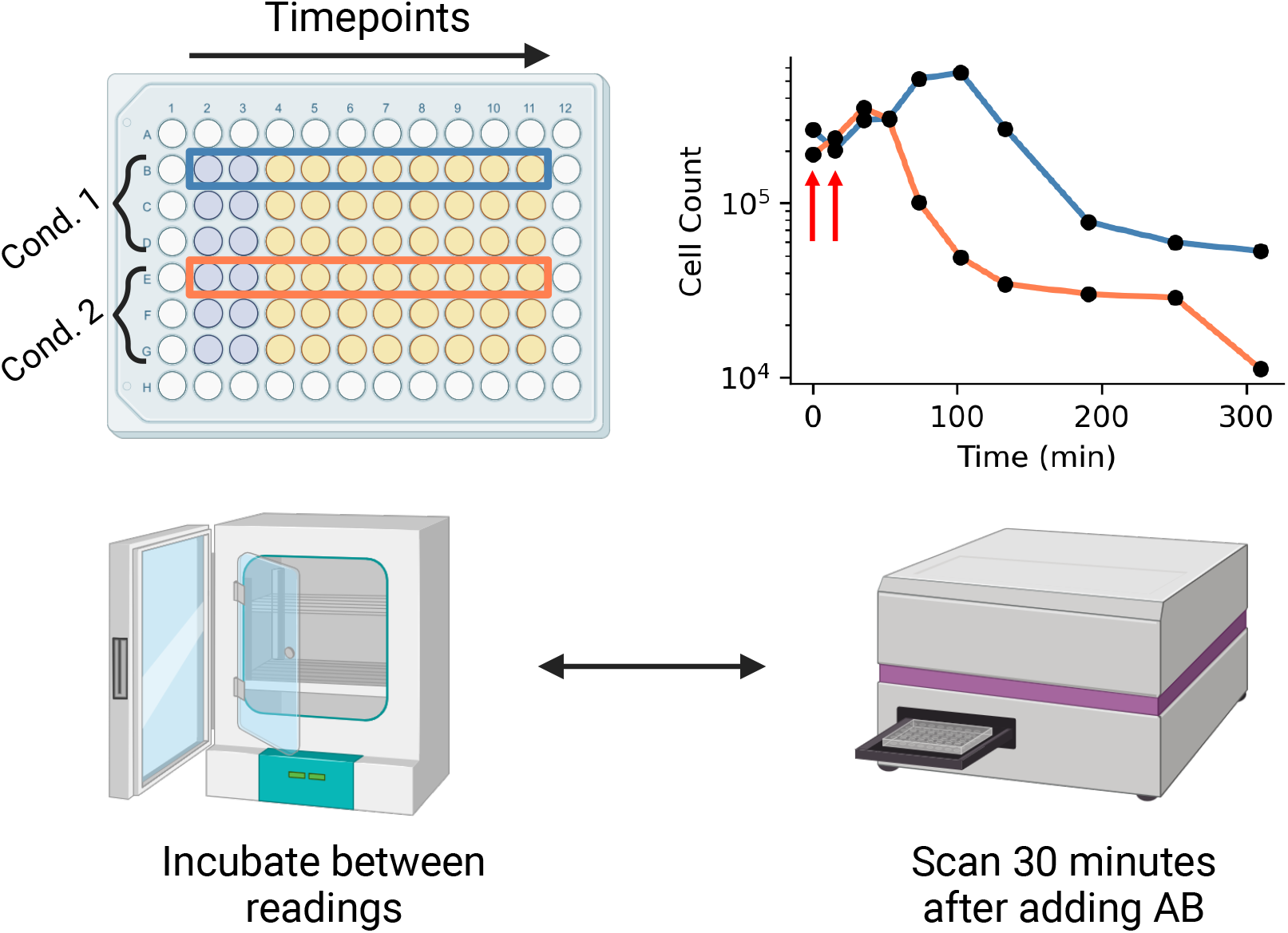
Time-kill assay overview. Cells are inoculated in 1 or more 96-well plates, with each plate containing 2 experimental conditions in triplicate (“Cond. 1” and “Cond. 2”). At the desired time point, resazurin is added to a column and the plate is scanned after 30 minutes. Cell count over time traces from rows B and E are shown on the right (10 and 100 µg/mL CTX concentration, respectively). In this illustration, AB has been added to columns 2 and 3, corresponding to timepoints 1 and 2 in the cell count over time plot (illustrated with red arrows). This process is repeated at the desired timepoints for each of the 10 rows (rows 1 and 12 in the plate are excluded). At the end of the experiment, for each row we can obtain an estimate of cell count over time.

Our proposed method improves upon previous time-kill assays in several ways. First, we show that time-kill estimates can be performed in triplicate for 6 different drug concentrations using only three 96 well plates, dramatically reducing the footprint relative to a comparable CFU-based assay. The method is also easily carried out by a single researcher in a single day, without the need for continuous bench work. In addition, our method does not rely on cell culture regrowth and thus is not biased by drug carried over from the experimental condition to the cell count assay; by incubating for only 30 minutes, our method reports a “snapshot” of the viable cell count at a single point in time. Finally, we show that our proposed method is robust to cell aggregation triggered by antibiotic exposure, which may prevent uniform sampling required for accurate CFU assays^25^.

## Materials and Methods

### Experimental model

We used *Escherichia coli* DH5*α* carrying the cloning vector pBR322 provided by the Weinreich Lab at Brown University^26^. Cells were cultured overnight from frozen glycerol stock in Luria-Bertani (LB) broth and 10 µg/mL tetracycline (MP Biomedicals).

### Key equipment

For fluorescence and optical density scanning, we used a Tecan Spark multimode microplate reader. Settings used for all experiments are shown in **Table 1**. The optimal gain for AB fluorescence scans may need to be adjusted for different experiments, in which case a new calibration curve will be required.

**Table 1.** Tecan settings for fluorescence and OD_600_ scans.

|  | Fluorescence setting | OD setting |
| --- | --- | --- |
| Shaking duration (s) | 10 | 10 |
| Shaking frequency (rpm) | 510 | 510 |
| Mode | Fluorescence Top Reading | Absorbance |
| Measurement wavelength (nm) | n/a | 600 |
| Excitation wavelength (nm) | 560 | n/a |
| Excitation bandwidth (nm) | 20 | n/a |
| Emission wavelength (nm) | 590 | n/a |
| Emission bandwidth (nm) | 20 | n/a |
| Gain | 40 | n/a |
| Mirror | Automatic (50% Mirror) | n/a |
| Number of flashes | 30 | 10 |
| Integration time (μs) | 40 | n/a |
| Lag time (μs) | 0 | n/a |
| Settle time (ms) | 0 | 50 |
| Z-position (μm) | 20000 | n/a |
| Z-Position mode | Manual | n/a |

### Method details

#### Generating calibration curves

We generated a curve to estimate bacterial cell count from AB fluorescence using a two-step procedure (**Fig 3**). First, we generated a cell count to optical density calibration curve. *E. coli* cells were inoculated from frozen glycerol stock in 3 mL LB broth and incubated overnight at 37° C and 220 rpm with the cap in the “vent” condition. We made a series of 2-fold dilutions of the culture in fresh media using 90 µL cell culture per well in a clear, polystyrene 96-well plate, with rows B-G serving as technical replicates and columns 2-10 comprising the dilution gradient. Column 11 was used for background estimation by adding fresh medium only. Optical density was measured using a microplate reader, and samples from row B, columns 2, 4, 6, and 8 were subjected to a colony forming unit (CFU) assay to quantify cell count. These samples corresponded to 1-, 4-, 16-, and 64-fold dilutions from the initial culture. We diluted each sample by 100,000-fold and plated 50 µL onto agar plates in triplicate. Note that different dilutions may be necessary for different organisms. Colonies were manually counted after overnight incubation at 37° C. We then background-subtracted the optical density data and fit a linear model to the log-transformed optical density and cell count data. We used the slope and y-intercept to estimate cell count from optical density using a linear fit of the log-transformed cell count and OD_600_ values.

**Figure 3.**
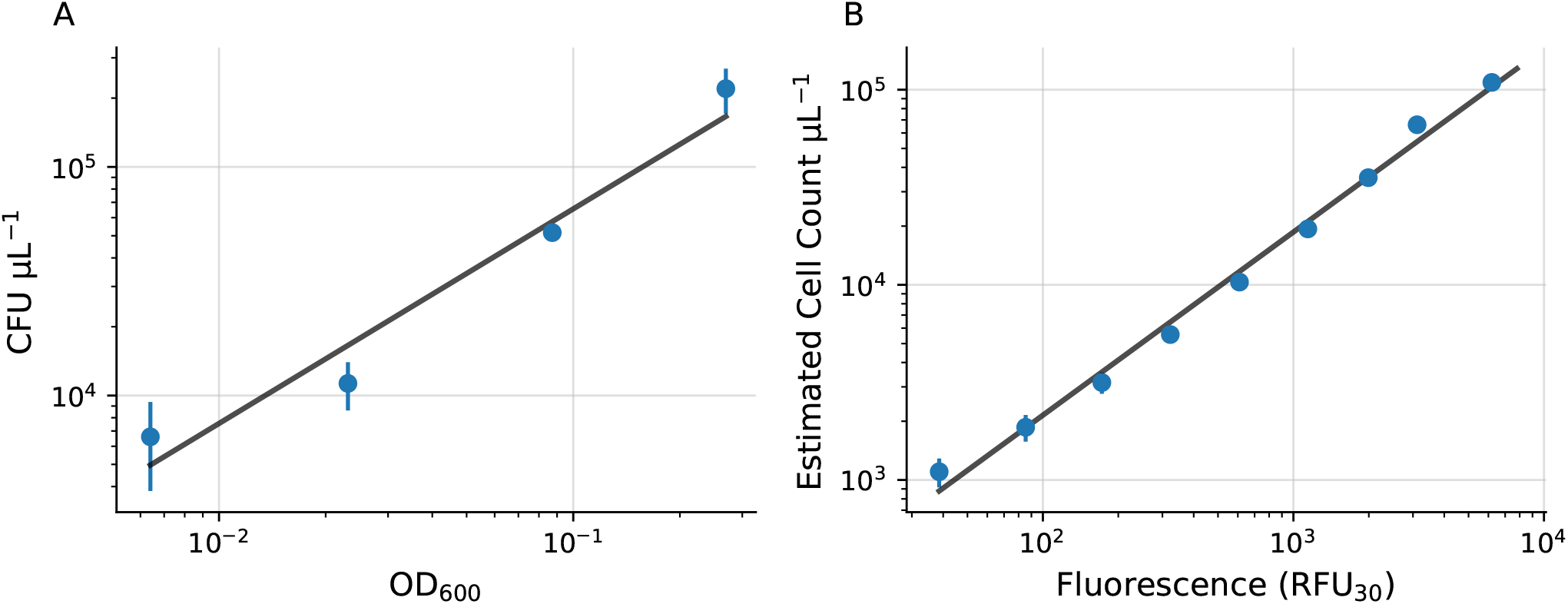
Optical density and fluorescence to cell count calibration curves. **(A)** Cell count versus optical density (OD600) and estimated linear fit in log-space. **(B)** Cell count versus fluorescence (RFU_30_) with linear fit in log-space. Error bars represent standard error of the mean. Some error bars are smaller than the plotted data point.

To estimate fluorescence, 10 µL AB was added to each well of the same 96-well plate (excluding row B used for the CFU assay) and the plate was incubated for 30 minutes at 37° C with 220 rpm shaking. After incubation, fluorescence was measured with a microplate scanner (excitation filter 540 nm, emission filter 590 nm). After background subtracting, we used the optical density-CFU calibration curve above to estimate cell count as a function of relative fluorescence units (RFU_30_). As before, we estimated the calibration curve with a linear regression on the log-transformed data.

#### Estimation of dose-response curves with time-kill assay

Three 96 well plates were prepared with each plate containing 2 conditions and 3 experimental replicates per condition. Rows B-G were used for experimental replicates while columns 2-11 were used for different timepoints. To optimize the time each condition spent in the dynamic range of the assay, we used different starting concentrations of cells for the two above- and below-MIC conditions. For the above-MIC conditions (conditions where the cell count was not expected to increase), we initialized the experiment with ∼2x10^5^ cells/mL, which we obtained by diluting the overnight culture by 4x with fresh media. For the below-MIC conditions, we initialized the experiment with ∼2 *∗* 10^3^ cells/mL by diluting the overnight culture by 100x. All experiments were initialized after 1 hr of pre-incubation after diluting the overnight culture. We prepared a 20,000 µg/mL stock solution of CTX following the manuscturer’s instructions. To ensure that each condition was exposed to drug at roughly the same time, 10 µL of the CTX solution (Thermo) was added to the wells for each condition first, followed by 90 µL of cell culture. The wells on the outer boundary of the plate were left empty, as fluorescence readings were found to be unreliable for these wells.

At time 0, 10 µL AB was added to column 2 of each plate, and plates were incubated for 30 minutes at 37° C with 220 rpm shaking. After 30 minutes, fluorescence was measured with the microplate scanner. 10 µL of AB was added to column three 15 minutes after adding AB to column 2, and plates were incubated for 30 minutes before performing the corresponding fluorescence scan. This process was repeated for columns 4-11, with plates being scanned 30 minutes after adding AB to each column. We used 15-minute time intervals between adding the AB in columns 2-6, 30 minute time intervals for columns 6-8, and 1 hour time intervals for columns 8-11. This approach resulted in sampling time points of roughly 0, 15, 30, 45, 60, 90, 120, 180, 240, and 300 minutes. This sampling scheme allowed us to observe the short- and long-term dynamics of the change in cell count over time. We applied the RFU_30_-cell count calibration curve from above to estimate the cell count at each time point.

For the cell aggregation experiments treated with DNase, 10 µL of cell culture was added to 90 µL of DNase solution comprising of 10 µL reconstituted rDNase and 90 µL DNase reaction buffer (Machery-Nagel). This resulted in a final DNase concentration of ∼0.5 U/µL. Samples were incubated for 15 minutes before depositing onto agar plates.

#### Net Growth Rate Estimation

Estimating the parameter of interest from time-kill data is a challenge, especially during cell death. While others have used parametric equations constructed from first principles^12^, we found overfitting to be an impassable challenge. Furthermore, we observed a variable death rate in our high drug concentration conditions. As such, a simple model with fewer parameters may be a more robust approach. As an illustration, we sought to estimate the maximum rate of change in cells using a linear regression on the log-transformed cell count data. To choose a subset of the data that balanced linearity with including as much data as possible, we defined the following objective function:

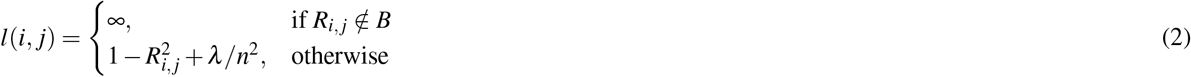

where *i* and *j* are the start and end indices for the subset, *R*_*i, j*_ is the correlation coefficient for the subset, 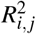 is the coefficient of determination, *n* is the number of points included in the subset, *λ* is a regularization parameter, and *B* is the boundary-defining set. Since 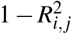 is strictly between 0 and 1, we set *λ* equal to 1. We also defined a boundary set *B* for the estimated slope of the regression based on whether the cell count was increasing or decreasing. For instance, for the drug-free condition, the slope must be greater than 0 since the cell count increased, so *B* = *{x* : *x >* 0*}* . If *R*_*i, j*_ is not in the set *B*, then the loss function is set to infinity. We then minimized *l*(*i, j*) for each dataset by an exhaustive search of all subsets larger than 2 elements. For the optimization step, time and cell count for each dataset were normalized between 0 and 1. We then used the subset identified with this process to estimate the maximum rate of change in cell count using a linear regression.

We compared this novel method (“optimized range”) to three other quantification techniques: “naive” linear regression”, “reduced range” linear regression, and area under the curve (AUC). For the naive linear regression, we simply fit a linear regression to the entire log-transformed dataset for each condition. For the reduced range linear regression, the data range used depended on whether or not the net change in cells was positive or negative by the end of the experiment. If the net change was positive (i.e. in the sub-MIC conditions), the data range used included the first data point to the data point with the maximum cell count. If the net change was negative, the data range included the data point with the maximum cell count to the last data point. Finally, we computed the AUC with numerical integration using the trapezoidal rule and normalized the result to between -1 and 1.

All linear regressions, curve-fits, and AUC calculations were performed with the SciPy python package. All code and data required for reproduction is available in the github repository: https://github.com/eshanking/time-kill-protocol.

## Results

### Estimating cell count over time with a time-kill assay

We first generated a calibration curve to estimate *E. coli* cell count from AB fluorescence. To generate an OD_600_-cell count calibration curve, we measured the optical density of a set of serial cell culture dilutions and directly estimated cell count using a colony-forming unit (CFU) assay (**Fig 3A**). Then, we measured optical density and AB fluorescence (RFU_30_) on the same set of cell culture dilutions and applied the OD_600_-cell count calibration curve to generate a RFU_30_-cell count calibration curve (**Fig 3B**). For both experiments, we used a linear fit of the log-transformed data to estimate the calibration curves.

To demonstrate our novel time-kill assay, we quantified a dose-response curve in *E. coli* by measuring cell count over time for different drug concentrations. We used this data to estimate the net growth rate as a function of drug concentration. *E. coli* were exposed to CTX concentrations below, at, and above the reported MIC (0.088 µg/mL)^26^. We found that 3 plates, which allowed for 6 experimental conditions with 3 technical replicates each, were the maximum number of plates that one person could manage logistically during the experiment. For each plate, condition 1 occupied rows B-D, while condition 2 occupied rows E-G (**Fig. 2**). Each column of the plate represents a different amount of time that the cell culture has been exposed to the corresponding drug concentration before sampling.

Example results with linear fits for estimating the maximum rate of change in cell counts are shown in **Fig. 4**. As expected, there was rapid logistic growth under sub-MIC conditions (0 and 0.01 µg/mL). Near the MIC (0.1 and 1 µg/mL), there were relatively small changes in cell count over time, although there was a slight downward trend. At higher concentrations (10 and 100 µg/mL), there was rapid cell death. Interestingly, in the 10 and 100 µg/mL conditions, there appeared to be dose-dependent lag in the onset of the antibiotic effect – the rapid decrease in the number of cells occurs earlier in the experiment for the 100 µ/mL condition than the 10 µ/mL condition. This phenomenon has previously been observed in similar time-kill experiments^12^.

**Figure 4.**
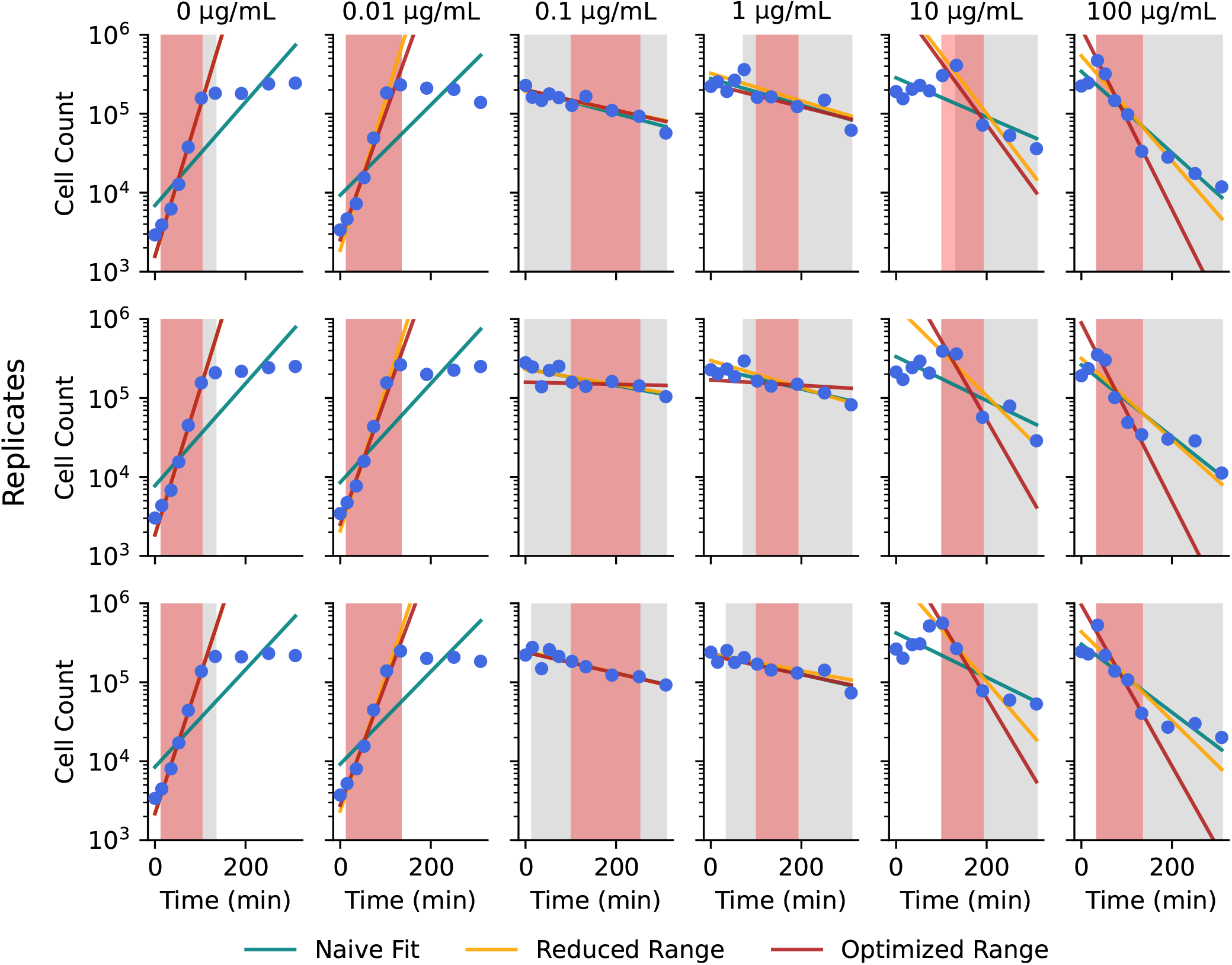
Cell count over time estimated from time-kill experiment. Each column represents a single drug condition with 3 experimental replicates. Columns are labeled at the top with the CTX concentration. Blue dots are estimated cell count values. Solid lines indicate different net growth rate estimates: ‘Naive Fit’ (linear fit to the entire data range), ‘Reduced Range’ (linear fit from either the start to the maximum or maximum to the end), and ‘Optimized Range’ (linear fit to the best estimated linear subset). Gray shading shows the range used for the Reduced Range fit, while red shading shows the range used for the Optimized Range fit.

### Pharmacodynamic curve estimation

As the sub-MIC conditions exhibit logistic growth, there are a variety of methods one may use to estimate the net growth rate, including calculating the slope of the exponential growth phase. This is also the case near the MIC, as the change in cells over time is roughly linear in log-space. However, the higher drug concentrations appear to exhibit non-constant net growth rates over time, with an initial small increase in cell count, followed by rapid decrease in cells, and concluding with a slight tapering off of the death rate. The precise parameter of interest to be estimated from the cell count data depends on the particular experimental question. As an illustration, we quantified the pharmacodynamic relationship using three different net growth rate estimates (naive, reduced range, and optimized range) and AUC (see methods) (**Fig. 5**). Data were fit to the following dose-response equation using SciPy,

**Figure 5.**
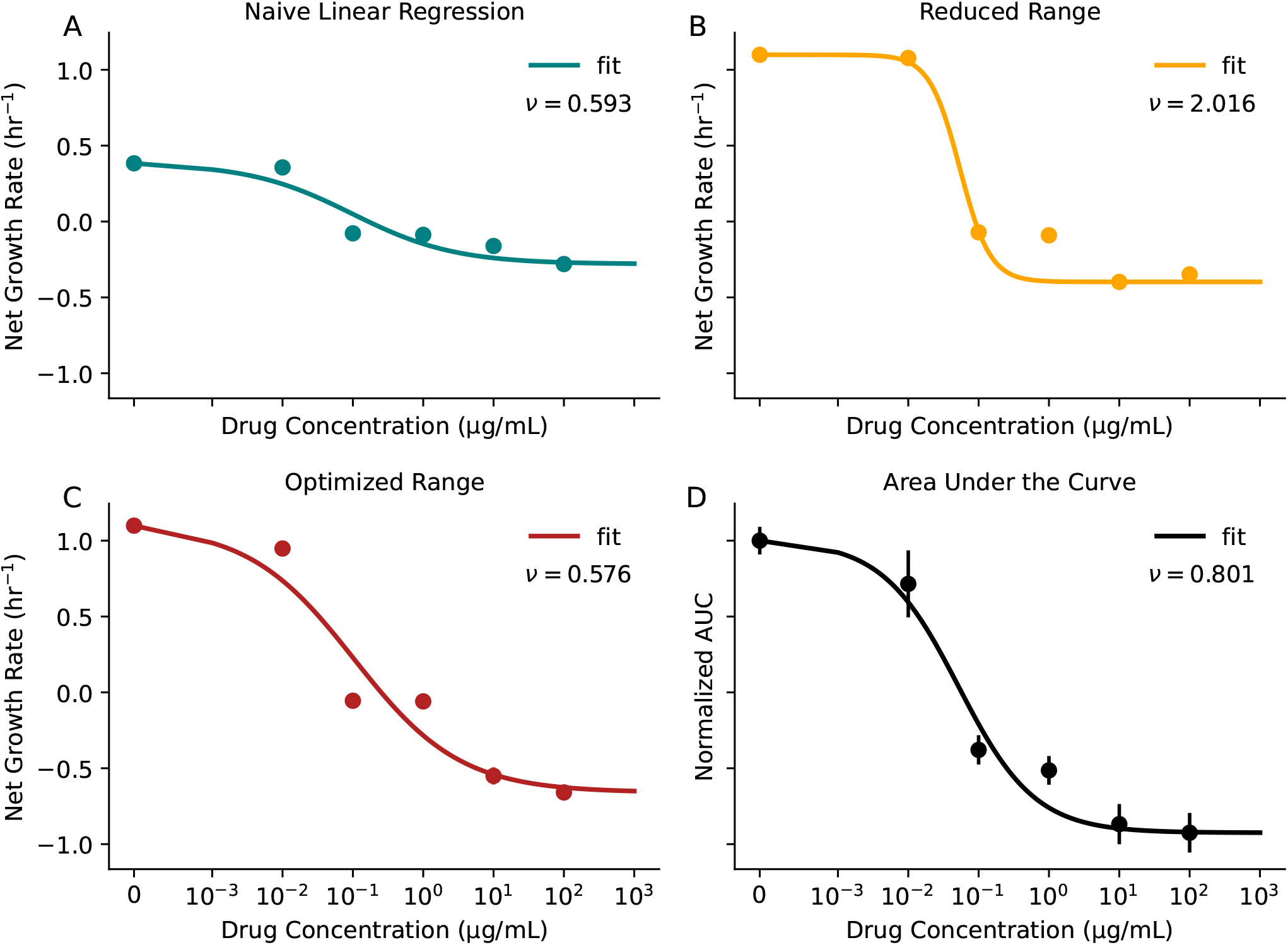
Pharmacodynamic curves for net growth rate estimates and AUC. **(A-C)** Pharmacodynamic curves with fits for different methods of net growth rate estimation. Colors correspond to the linear fits shown in **Fig 4. (D)** Normalized AUC versus drug concentration. Solid lines show the dose-response curve fits (**Eq 3**). The Hill coefficient *ν* for each curve is shown inset. Error bars represent standard deviation of the net growth rate or AUC estimate (in many cases, the error bar is smaller than the size of the plotted datapoint).

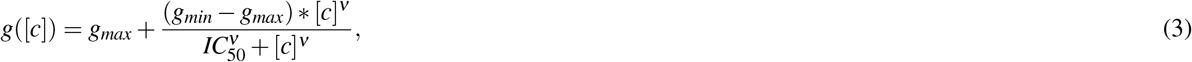

where [*c*] is the drug concentration, *g*_*max*_ is the maximum value (growth rate or AUC), *g*_*min*_ is the minimum value, *IC*_50_ is the half-maximal inhibitory concentration, and *ν* is the Hill coefficient, which describes the steepness of the curve. Importantly, all four methods of quantification reveal dose-dependent cell death, with curves that cross zero as the drug concentration increases. We also found that the AUC method may be more sensitive to small changes in net growth rate, as the AUC for the sub-MIC 0.01 µg/mL is slightly lower than the AUC for the no-drug condition (0.71 verus 1.0, respectively). In contrast, the growth rate estimates are roughly the same for both conditions (**Fig 5**).

### Comparison to OD in cytotoxic and cytostatic drugs

In order to fully understand the capabilities of our method compared to an OD-based approach, we simultaneously measured OD and AB fluorescence in a time-kill assay. We repeated this experiment in both CTX, which is a bactericidal drug, and tetracycline (TET), which is a bacteriostatic drug. Tetracycline inhibits bacterial growth by binding to the 30S ribosome subunit and arresting translation^27^. Despite being classified as bacteriostatic, tetracycline, in many cases, may result in cell death – this is because bacteriostatic drugs are defined as drugs whose minimum bactericidal concentration (MBC) is at least 4 times higher than the MIC^28^. MBC refers to the drug concentration that results in 1000-fold reduction in cell count after 24 hours. Therefore, a given drug can have substantial killing activity and still be classified as bacteriostatic.

In the CTX experiment, there were clear dose-dependent death dynamics and dose-dependent lag in the onset of the killing effect when measured with the AB assay (**Fig 6A**). However, the OD assay exhibited a steep drop off in detected cells after ∼150 minutes for the 10 and 100 µg/mL conditions. By visual observation, this drop off in OD was driven largely by cell aggregation, leaving portions of the well almost entirely clear of cells (representative microscopy shown in **Fig S1**). However, cell aggregation was not grossly apparent in the low drug conditions (0.1 and 1 µg/mL). As a result, OD measurements in the high-drug conditions do not reflect a true lack of viable cells – this is explored in detail in the following section. In addition, technical replicates in the AB assay were very similar, with small standard deviation, while technical replicates in the OD assay exhibit much higher variance. This can also be explained by cell clumping within the well causing unreliable readings. Furthermore, the AB assay revealed a gradual loss of cells after ∼150 minutes in the 0.1 and 1 µg/mL conditions, while the OD assay suggested an increase in cells. This discrepancy may be partially explained by dead cell debris obscuring the OD reading, as it is not clear how lysed or nonviable cells impact optical density.

**Figure 6.**
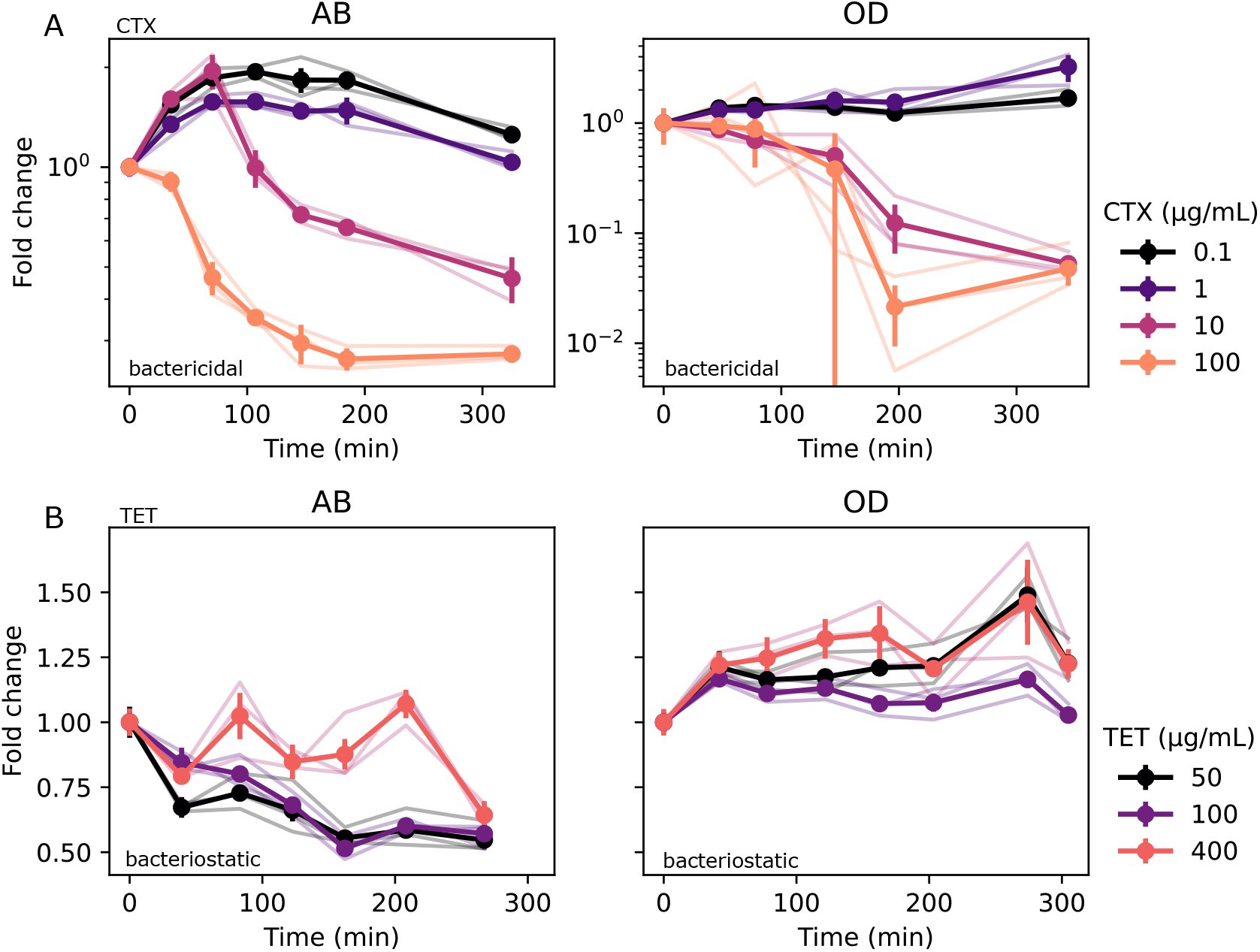
OD cannot resolve cell loss in cytostatic or cytotoxic drugs. **(A)** Time-kill assay in CTX measured with AB fluorescence (left) and OD (right). Light traces are individual replicates, which dark traces with error bars are means of the technical replicates. **(B)** Time-kill assay in TET measured with AB fluorescence (left) and OD (right). Error bars are standard deviation. *N* = 3 technical replicates for each measurement.

In the TET experiment, we similarly observed a decline in cell count with the AB assay, although there was no clear pattern in dose-dependent death rate or dose-dependent lag (**Fig 6B**). In contrast, the OD assay suggested a relatively flat or gradual increase in cell count over time. As observed in the previous experiment, this discrepancy may be explained by dead cell debris impacting OD estimates.

### Cell aggregation biases CFU assay

In order to validate the estimated cell count from the fluorescence assay, we performed a standard colony-forming unit (CFU) assay in parallel for 0 and 10 µg/mL CTX (**Fig S2**). For CFU estimates, samples were diluted 10^4^ and 10^5^ times and plated on antibiotic-free LB agar plates. Individual colonies were then counted by hand after 24 hrs incubation. While AB and CFU cell count estimates for the no-drug condition aligned closely, the CFU estimate for the 10 µg/mL CTX condition diverged sharply after the initial measurement (CFU = *∼* 3 *∗* 10^3^ cells µL^-1^, AB = *∼* 1.8 *∗* 10^5^ cells µL^-1^ after 80 minutes).

In all iterations of the time-kill experiment, we observed significant visible cell aggregation within wells with drug (**Fig S1**) – this may explain the difference between the two estimates, as cell aggregation may prevent uniform sampling of cells during the CFU assay. Recent work has identified extracellular DNA (eDNA) as a driver of cell aggregation in *E. coli* when exposed to drug^25^. To explore this further, we repeated the CFU experiment with and without recombinant DNase. During sampling, we incubated 10 µL of cell culture with 90 µL DNase solution or control for 15 minutes. We visually observed a rapid loss of the cell aggregate in the DNase condition, while the aggregate in the control condition remained visible. CFU quantification revealed significantly higher cell counts in the +DNase condition after 180 minutes of exposure to drug, demonstrating that cell aggregation significantly reduces the CFU assay cell count estimate (**Fig 7A**).

**Figure 7.**
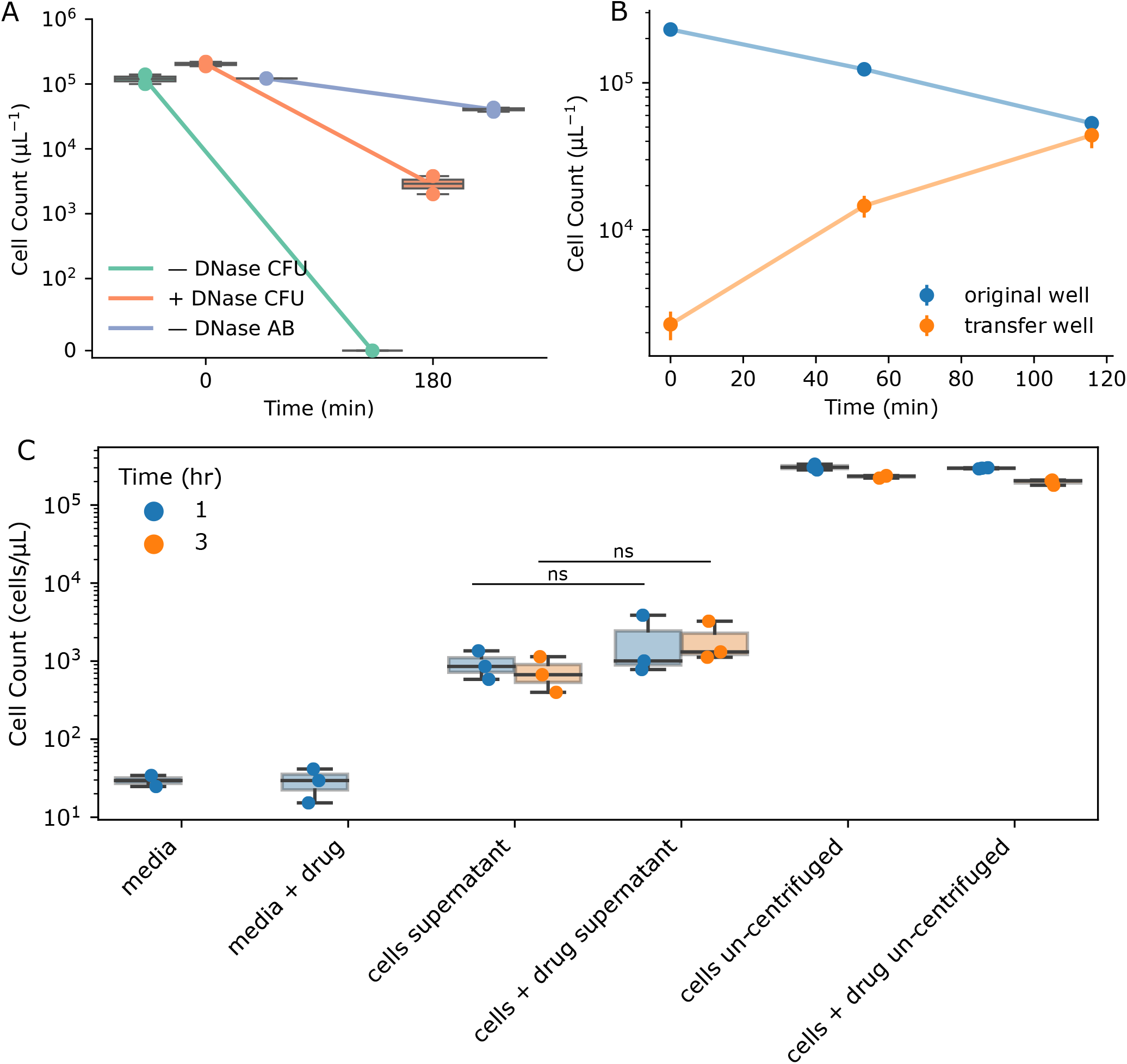
Investigating the impact of cell clumping and cell lysate. **(A)** Estimated cell count for two timepoints under 3 conditions: +DNase, –DNase, and AB fluorescence. +/– DNase conditions were estimated with direct CFU assay. CTX concentration = 10 µg/mL for each condition. **(B)** Cell count estimated from AB fluorescence over time while transferring the visible cell aggregate from the ‘original well’ to the ‘transfer well’ containing fresh media. CTX concentration = 10 µg/mL in the original well. **(C)** Estimated cell count by sampling either directly from cell culture (cells un-centrifuged) or from centrifuged supernatant (cells supernatant). Blue color indicates a 1 hr sample time while orange indicates a 3 hr sample time. CTX concentration = 10 µg/mL for the + drug conditions. Error bars represent standard deviation. *N* = 3 technical replicates for each experiment.

Based on these results, we hypothesized that cell aggregates contain viable cells that can reduce resazurin leading to measurable fluorescence signals, and these viable cells are not detected with a traditional CFU assay. To investigate this further and quantify the time dynamics of cell aggregation, we repeated our time-kill assay while manually transferring the visible cell aggregate from wells with 10 µg/mL CTX to wells with fresh media. At the designated sample time, we used a pipette tip to scoop the aggregate from the cell culture well and deposit it into a well with 90 µL fresh media (“transfer well”), after which we added 10 µL AB. We also added AB to the original well from which the aggregated was sampled from (“original well”). At the 0 time point, we simply dipped the pipette tip in the original well and the transfer well, as no visible aggregates had formed. We observed a time-dependent increase in estimated cell count from the transfer well along with a corresponding decrease in estimated cell count from the original well (**Fig 7B**). These time dynamics show that the number of viable cells in the aggregate increases over time after exposure to drug while the number of free-floating cells decreases over time.

### Dead cell debris does not impact cell count estimate

To determine whether non-viable cells may bias the estimated cell count, we next investigated whether cell lysate significantly contributes to the fluorescence signal. We treated *E. coli* cell cultures for 1 hr and 3 hrs with either 10 µg/mL CTX or media control in 1.5 mL centrifuge tubes. At the designated timepoints, we centrifuged the samples, collected the supernatant, transferred 90 µL to a 96-well plate with 10 µL AB, and incubated for 30 minutes. After scanning, we applied the cell count-fluorescence calibration curve described above. We found no significant difference in the estimated cell count from the centrifuged control (cells supernatant) and centrifuged with drug (cells + drug supernatant) conditions (**Fig 7C**). These results suggest that, while extracellular material may contribute some fluorescence signal, this background signal is not driven by drug-mediated cell death. We also observed no time-dependence in the signal from either condition, further establishing that cell lysate does not significantly contribute to fluorescence. In addition, the estimated cell count from the supernatant was ∼2 orders of magnitude less than the estimated cell count from the un-centrifuged conditions; therefore, extracellular material cannot explain the difference between the cell count estimated from AB fluorescence and the direct CFU assay cell count from **Fig 7A**.

## Discussion

Here, we present a novel fluorescence-based time-kill assay in bacteria for estimating dose-dependent cell death rates. Our method improves upon previous techniques by dramatically reducing the material cost and experimental workload while being robust to drug carry-over and cell aggregation. Furthermore, the proposed method does not require genetic engineering or labeling of the model organism. Our method allows for the quantification of 6 drug conditions and 10 timepoints using only three 96 well plates and can be comfortably executed by a single researcher. We demonstrated our proposed method by quantifying dose-response curves in *E. coli* subject to cefotaxime.

While direct validation of cell count over time remains a challenge, our results suggest that (1) AB fluorescence reports a realistic cell count within a drug-induced bacterial aggregate and (2) extracellular debris from non-viable cells does not significantly impact the cell count estimate. While the cell count estimates for the + DNase CFU assay and the AB assay diverge significantly, there are several possible explanations – for instance, AB fluorescence reports the near-instantaneous number of viable cells, whereas CFU estimates rely on the regrowth of cells after overnight incubation. There may be viable cells that can metabolize resazurin that are nonetheless unable for form colonies due to drug exposure. Furthermore, although cell cultures are diluted for the CFU assay, there is still residual drug plated that may inhibit colony formation. Finally, DNase treatment may not have eliminated all aggregation, preventing uniform sampling and resulting in under-counting of viable cells.

While being a substantial improvement over previous time-kill methods, our proposed technique has several limitations. First, each timepoint requires intervention by a researcher, demanding attention over several hours. In addition, the dynamic range of the assay is limited, and cell death and cell growth conditions may need to be initialized at different starting cell densities to optimize time spent in the dynamic range. In this case, researchers should consider density-dependent effects that may bias comparisons between experimental conditions. Additionally, *a priori* knowledge of the model organism MIC is useful for determining initial densities. Finally, while not unique to the proposed method, estimation of the parameter of interest (such as net growth rate) remains a challenge. However, we provide an optimization method for linear range estimation.

This method opens the door for more detailed PK-PD modeling of clinical bacterial infections. By quantifying more precisely how different organisms respond to drug, we may better understand why certain clinical regimens fail to eliminate an infection. In addition, we may be able to more accurately model pathogen evolution within a patient, allowing us to predict evolution and optimize drug dosing regimens.

## Acknowledgements

JGS and ESK were supported by NIH 5R37CA244613-04 (https://www.cancer.gov/). ESK was supported by NIH 3T32GM007250-46S1 (https://www.nigms.nih.gov/). JGS was supported by American Cancer Society Research Scholar Grant RSG-20-096-01 (https://www.cancer.org/).

## Supplementary information

### Cell aggregation microscopy

Cell aggregation was grossly visible by eye in the 96 well plates when exposed to drug. Microscopy images revealed filamentous structures ∼500 µm in width (**Fig S1**). These structures made viable cell count estimates with optical density unreliable.

**Figure S1.**
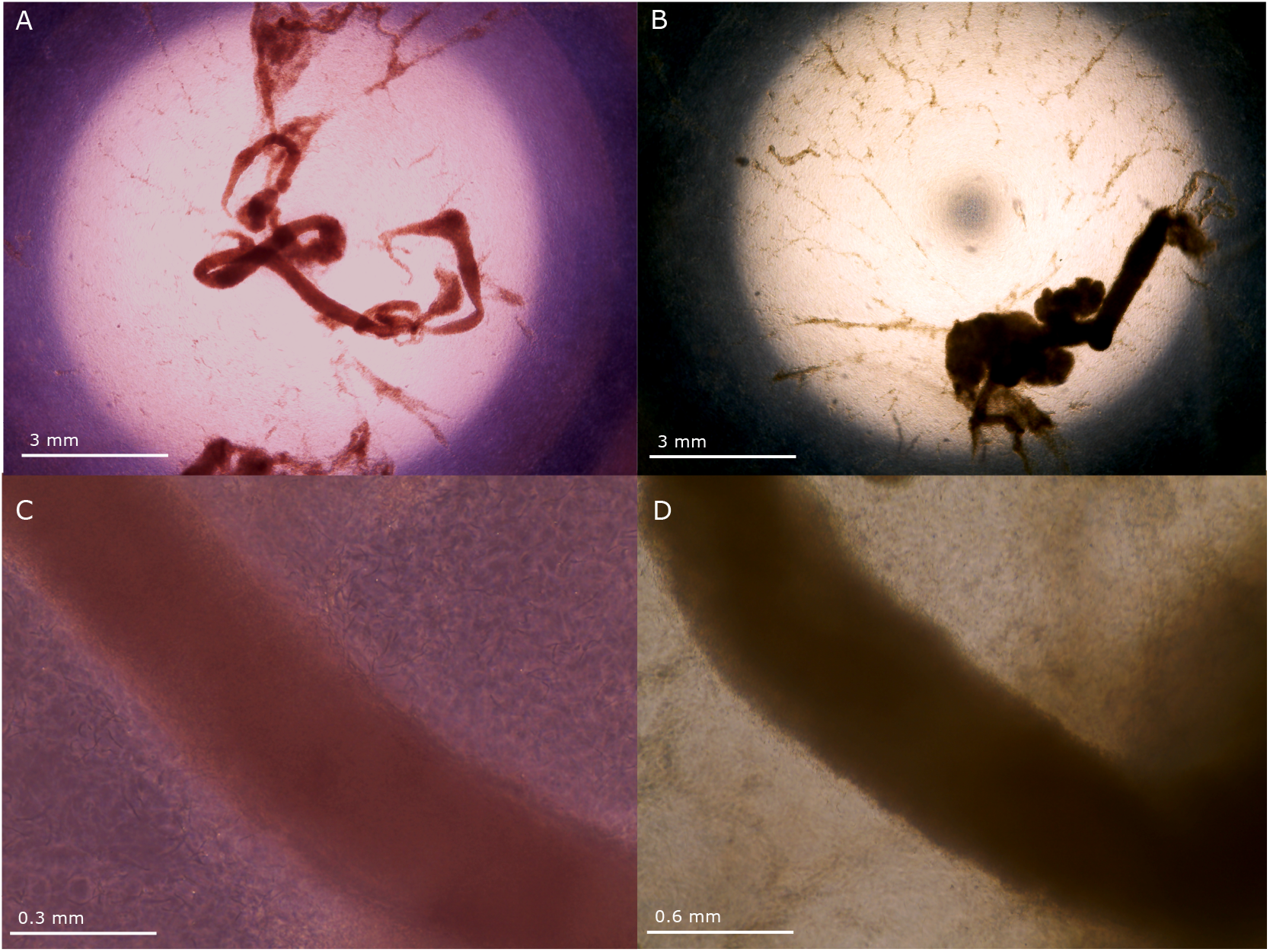
Representative microscopy showing E coli aggregation due to drug exposure. (A) 4 x magnification with AB added. (B) 4 x magnification with no AB (media only). (C) 40 x magnification with AB added. (D) 20 x magnification with media only.

### CFU assay validation

To validate the protocol, we first compared the cell count estimated from fluorescence to cell count estimated with a CFU assay. While CFU and AB estimates from the no-drug condition aligned closely, we found that estimates for the 10 µg/mL conditions diverged significantly after the first timepoint (**Fig S2**).

**Figure S2.**
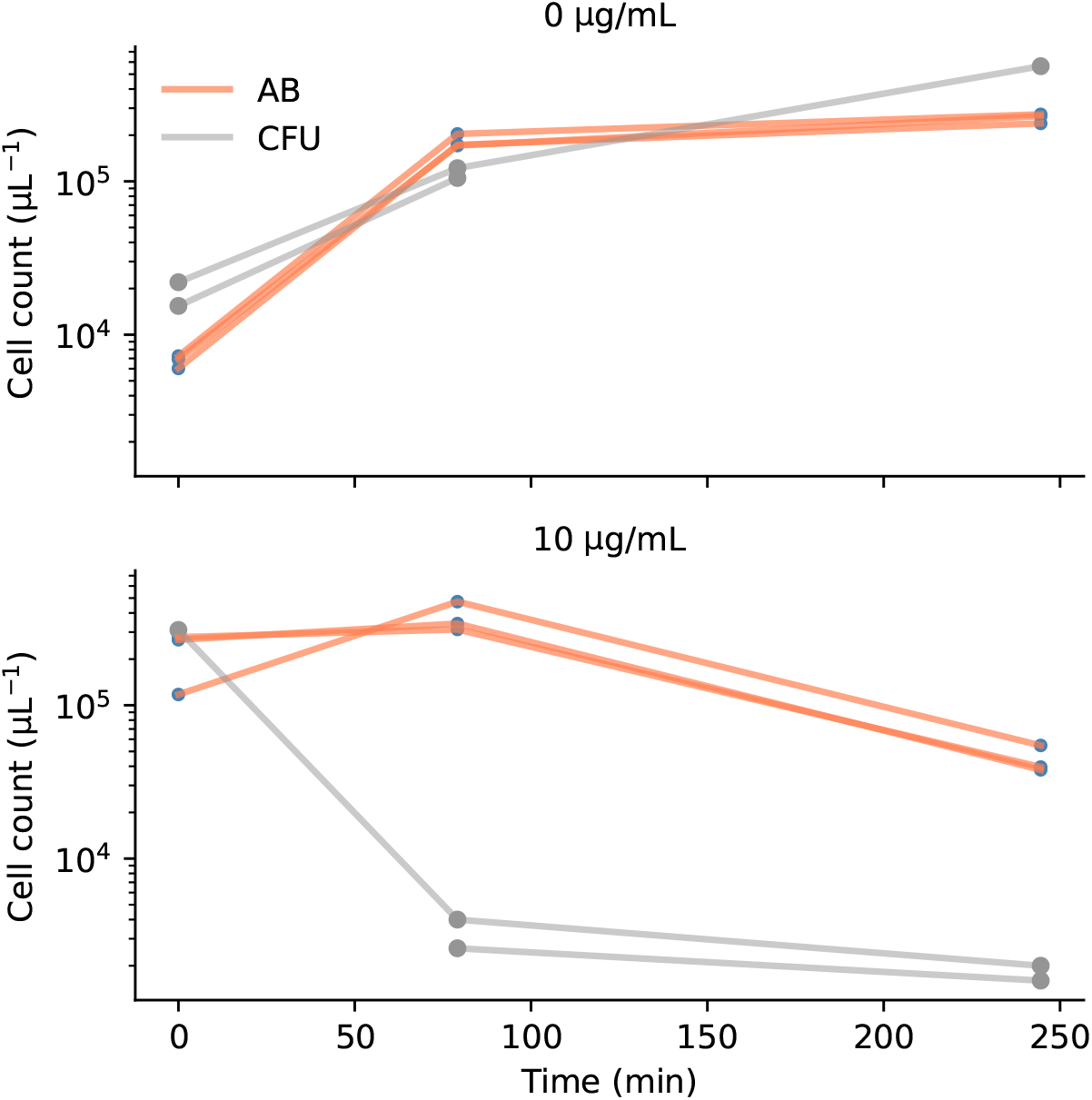
Comparison of cell count estimate with AB fluorescence and CFU assay. Top: no drug condition. Bottom: 10 µg/mL condition.

## Notes

https://github.com/eshanking/time-kill-protocol

### Competing Interest Statement

The authors have declared no competing interest.

### Summary of Updates

Added additional time-kill experiments in cefotaxime and tetracycline in the results section. Added microscopy data.

